# Preventing pathogenic dimerization in a misfolded antibody light chain through the design of an inhibitory peptide

**DOI:** 10.1101/2025.09.16.676588

**Authors:** Fausta Desantis, Mattia Miotto, Edoardo Milanetti, Giancarlo Ruocco, Lorenzo Di Rienzo

## Abstract

Immunoglobulin light chain (AL) amyloidosis is the most common form of systemic amyloidosis. The disease correlates with the formation of insoluble aggregates mostly composed by patient-specific antibody light chains, whose hypervariable regions make each case unique and highlight the need for personalized therapeutics. In this study, we focused on a pathogenic homodimer we previously obtained from a patient-derived light chain. By analyzing the dynamics and the interface of this dimer, we identified a 15-residue peptide with potential inhibitory activity. The peptide was then refined using a computational mutagenesis protocol that iteratively improved its sequence to maximize complementarity with the protein interface, taking into account shape, electrostatics, and hydropathy. The resulting optimized peptide is found to bind the monomer with a binding affinity comparable to that of the full pathogenic interface. These results suggest that the designed peptide could act as an effective antagonist of the pathogenic dimer, and demonstrate that our computational strategy could provide a general framework for designing patient-specific inhibitory peptides against aggregation-prone proteins.

## I. INTRODUCTION

Amyloidoses are a group of disorders characterized by the formation and deposition of insoluble protein assemblies with disabling and fatal effects [1]. Depending on the type and heterogeneity of tissues involved, they can be discriminated into localised amyloidoses, which have only one target (usually the central nervous system, as in the case of *Alzheimer’s disease, Amyotrophic Lateral Sclerosis* or ALS, *Parkinson’s disease*), and systemic amyloidoses, where multiple tissues can be affected, like *wild type Transthyretin* (ATTR), *Serum Amyloid A* (AA) and *Antibody Light Chain* (AL) amyloidosis [2, 3].

Although the association mechanism can vary depending on the molecular species involved, in most cases, aggregation is triggered by protein conformational changes [4]. The change in the free energy landscape underlying this protein misfolding is often caused by amino acid mutations which, by altering non-bonded intramolecular interactions, favor conformational states different from the physiological one [5]. This aspect is particularly relevant in AL amyloidosis, where the aggregating species are immunoglobulin light chains that are overproduced by defective B cells. This feature, at the core of the efficacy of our immune system, besides playing a pivotal role in the manifestation of the above-mentioned conformational changes, makes AL amyloidosis strongly patient-specific [6].

Nevertheless, a common aggregation pathway has been proposed, which originates, during the *lag phase*, precisely from this misfolding event [2, 7]. This misfolded conformation is thought to represent the molecular basis for the formation of a pathological homodimer [8]. Unlike physiological LC dimers, which play a protective role against aggregation [9, 10], this dimer represents the precursors of the aggregation pathway. After this, a cascade of events follows, ultimately leading to the formation of amyloid fibrils that primarily affect the heart and kidney [2]. Interestingly, the amyloid fibrils associated with the insurgence of this condition have been demonstrated to be mostly composed of variable domains, *V*_*L*_ (although evidence exists of the presence of full-length light chains [11] or constant domains alone [12], as well).

In this panorama, disrupting the association from the earliest stages could be a strategy against disease development. Along this line, we propose a method to design a molecule apt at impeding dimerisation in AL amyloidosis, a case study system that we had previously addressed [13]. A common procedure for the treatment of AL amyloidosis is chemotherapy-induced suppression of the amyloid synthesis by targeting the B cell. However, the extant drugs relying on antibodies and small molecules aimed at inhibiting the aggregation of amyloid precursors or at dissociating already-deposited amyloid fibrils are still at a clinical evaluation stage [2] [14].

The workflow necessary from the detection of a suitable drug to its circulation is not trivial. Employing *in-silico* methods to model potentially efficient molecules may help shorten this path by moderating experimental development costs and real-world testing risks [15].

In this scenario, investigating the properties of the amyloid precursors and the consequent design of drugs based on binding-antagonist peptides may come to aid in the perspective of synergistic therapies. Indeed, peptides are involved in 40% of protein-protein interactions mediating a wide range of processes spanning from signalling to protein trafficking to immunology [16] and have proved to be efficient drug candidates [17] [18] thanks to their biological compatibility, low toxicity and specificity [19].

In a previous work [13], we performed extensive molecular dynamics simulations on the experimental structure of a patient-derived amyloidogenic mutant light chain [20] [7]. The simulations spotlighted a misfolding event characterized by the exposition of hydrophobic patches. Based on the misfolded configuration observed, we were also able to effectively predict the structure of the pathogenic homodimer, seed of the higher order aggregation. The aim of this work is to design a peptide capable of acting as a competitive binder to block the pathological binding interface and thereby prevent the subsequent aggregation process.

One of the most common and effective strategies in peptide design is based on extracting a sufficiently long linear segment from a protein–protein interface [21], so that it can function as an inhibitor. For this reason, the first step of our study was to characterize, through extensive molecular dynamics simulations, the pathological dimer we had previously identified. Hence, we identified a 15-residue-long peptide whose suitability is assessed in terms of residue occurrence at the interface and intermolecular energies.

To make this peptide effective in preventing homodimer association, it needs to be optimized. Indeed, since the peptide represents only a portion of the entire binding site, it is unlikely to achieve, energetically, the same binding gain as the full protein–protein interaction. We then move to optimize the peptide using stochastic mutagenesis protocol, based on a Monte Carlo algorithm, where the complementarity between the peptide and the protein is assessed using chemical-physical properties of the two interfaces. Specifically, the shape complementarity is measured by the application of the 2D Zernike formalism, obtaining a compact representation of the local shape of molecular surfaces [22].In addition, electrostatic complementarity is evaluated using a coarse-grained atomistic approach that calculates the Coulomb potential energy of the interface configuration [23]. Finally, a term accounting for chemical compatibility was also included, considering hydrophobicity based on an amino acid hydrophobicity scale previously developed by our group [24].

At each step of the Monte Carlo algorithm, a stochastic mutation was introduced in the peptide, and the complementarity between the target protein surface and the peptide was evaluated using the descriptors described above. The acceptance of each mutation was then determined according to a cost function that accounts for improvements in the interface based on these descriptors. It is worth noting that in recent years we developed similar design protocols that have proven highly effective both in inhibitory peptide engineering [25] and in various other protein optimization processes [26–29].

After the widespread application of the optimisation protocol to ensure an exhaustive sampling of the sequence space [30, 31], we selected the best peptide among the accepted sequences. We thus tested with extensive molecular dynamics simulation the obtained peptide to work as inhibitor molecule. As an external validation, the optimized peptide is remarkably characterized by a very high binding affinity toward the molecular partner, as predicted using the PRODIGY tool [32], comparable to the values obtained when the whole pathogenic dimer is considered. Hence, such investigations confirm the detected peptide as a suitable binding competitor and testifies efficiency of the mutagenesis workflow proposed.

## II. RESULTS AND DISCUSSION

### A. Search for a convenient initial peptide sequence

As a first step, we looked for an initial template for an antagonist peptide against pathological dimerisation. To do so, we scanned the dynamic features of the pathogenic dimer we obtained in [13]. Therefore, we performed a 1-*µ*s-long molecular dynamics of such a homodimer, aiming to identify the residues mostly involved in binding. Indeed, for each frame we identified the residues involved in contacting the molecular partner, the latter defined as all those residues whose distances between *Cα*s of the two protein partners are lower than 8 Å [33] [34]. We thus checked on the residues’ occurrence at the binding site along the simulation, reported in the barplot in Fig. 1a, which depicts the probability of each residue to establish a contact. One can see that two regions are mostly involved in binding, namely residues 47-62 and 77-84. In the context of identifying the longest linear segment within the binding site, it is important to note that residues in positions 47 to 62 are characterized by very high contact percentages, where the mean is 64% and none of them interact for a time lower than 10%.

**FIG. 1:**
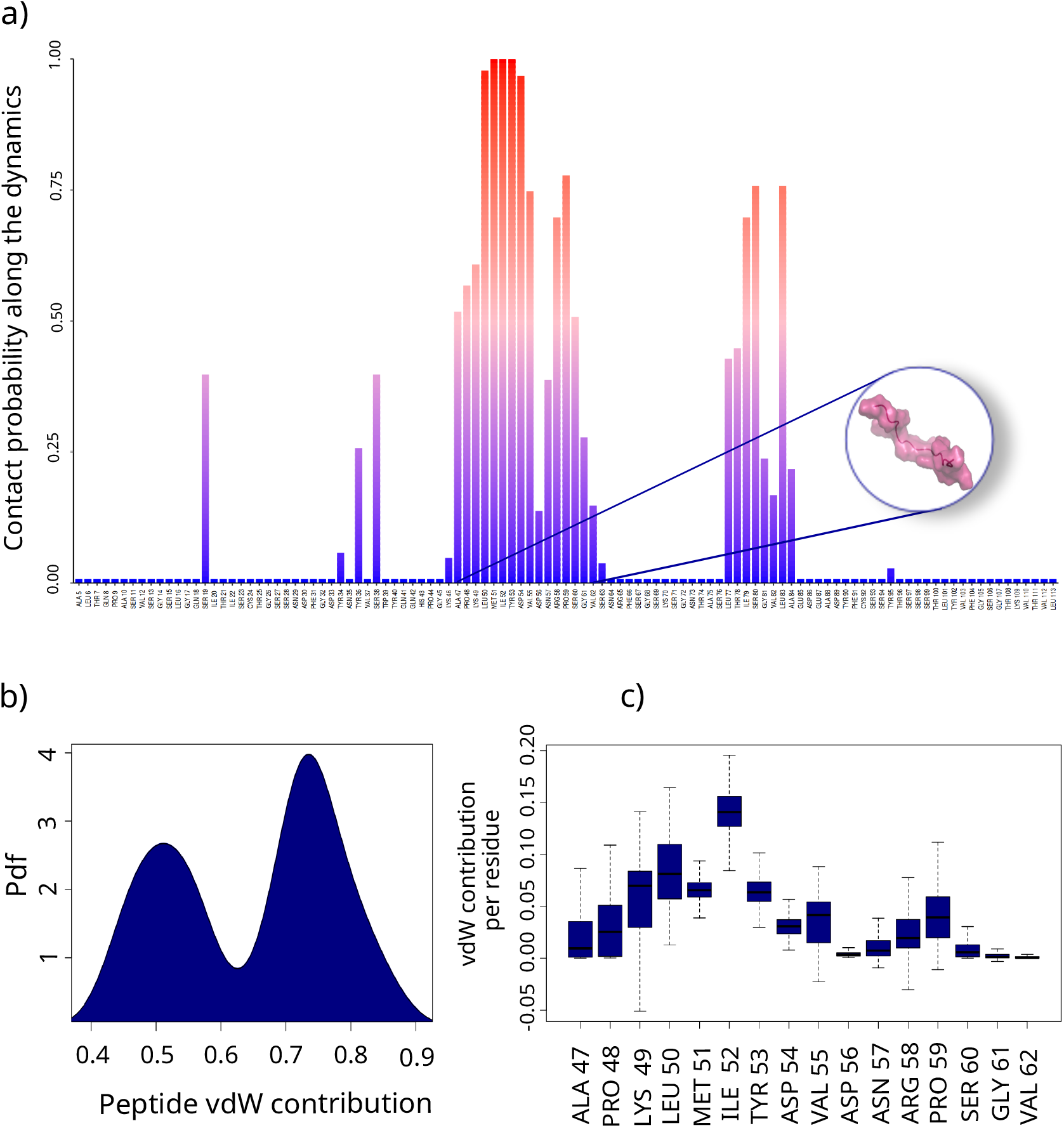
Docked interface analysis: **a)** Residue occurrence during the simulation time. **b)** Boxplot of the ratio between vdW energies of that residue and the rest of the interface for each residue in the peptide. **c)** Distribution of the global contribution of the peptide to the vdW interaction with respect to the rest of the interface.

To further assess the relevance of the selected peptide in complex stability, we appraised the relevance of such a region from an energetic point of view. Indeed, in a recent work, we evaluated the importance of the non-bonded energy network in determining the binding affinity in protein-protein complexes. We demonstrated that van der Waals (vdW) interaction energies at the interface are strongly correlated with binding affinity [35]. Therefore, for each frame we calculated the intermolecular vdW interactions between each pair of residues, aiming at evaluating the fraction of energies dependent on the analyzed portion. In Fig. 1b we report the probability distribution of such values, demonstrating that such part is very important for the dimer stability: this region is responsible on average of the 65% of the entire vdW interactions, where in each frame the whole is always responsible for more than 40% of the energy exchange, reaching in some case even 75% of the total energy.

Interestingly, looking at how this interaction is distributed among the amino acids of the peptide (see the boxplot in Fig. 1c), ILE 52 turns out to be the most interacting residue – contributing up to the 20% of the interaction – together with residues LYS 49, LEU 50, MET 51 and TYR 53 which participate in the interaction with a median value higher than 5%. Notably, residue 52 had already been reported as one of the two hydrophobic residues exposed upon misfolding, and as a key determinant in pathological dimer formation. The present finding further supports its importance in dimer stability [13]. Conversely, the C-terminal residues seem to be less important from this perspective.

Summarising, the above analysis suggests that the selected linear protein region can serve as a good template to work as a peptide binder to the misfolded light chain to impede pathogenic dimerisation.

As the optimisation procedure that will be exposed in the next section acts on the interface of the homodimer, a central assumption of this study is that, in solution, the peptide can fold into the same conformation observed at the protein–protein interface. Such structural consistency is necessary for the peptide and its optimized variants to act as effective inhibitors. Therefore, the conformational space explored by the peptide isolated in solution must be compatible with those that it spans when lying within the original pathogenic interface or when alone but bound to the chain. More specifically, the conformation the peptide adopts inside the protein-protein interface must also be maintained when isolated.

To examine whether such requirements are fulfilled, in addition to the dimer simulation we performed a 1*µ*s-long molecular dynamics of the peptide both in isolation and when bound with the other chain of the pathogenic dimer. The results of the simulations in the three cases ((i) peptide belongs to the original interface, (ii) isolated in solution, (iii) bound to the molecular partner) are depicted in Figure 2. Looking at the Root Mean Square Deviation (RMSD) (left panels in Fig. 2), as expected the most stable is the one where the peptide behavior is derived directly from the belonging interface, while the largest conformational changes are achieved when the peptide is isolated in solution. On the other hand, although the mobility of the peptide increases when it is bound to the other chain without the presence of other constraining interface neighbours, its conformation still appears to be quite stable and compatible with the case in which it belongs to the protein-protein configuration.

**FIG. 2:**
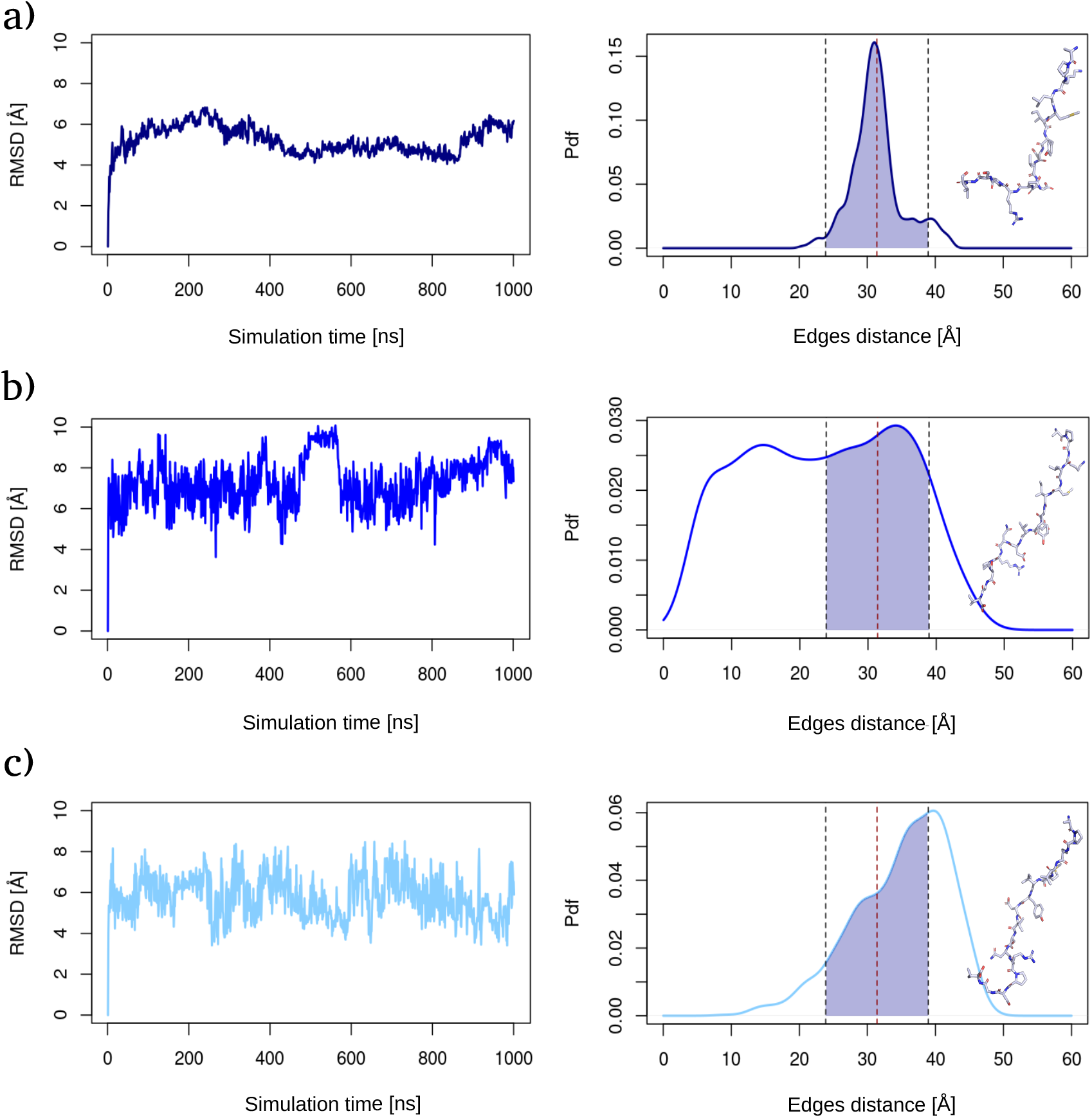
Dynamical properties of the peptide: **a)-c) left panels** RMSD of the peptide when belonging to the original interface (navy blue), alone (blue) and when bound to the other chain (light blue). **a)-c) right panels** Terminal C-*α*s distances distribution, when bound to the other chain, for the peptide in isolation. The blue-shaded area in d) refers to the portion of dynamics in which the peptide assumes an extended conformation (92.11% of the simulation). The shaded areas in b) and c) highlight the portion of the distribution corresponding to the extension of the peptide as depicted in a) and represent 58.94% and 42.11% of the respective simulations. Each box also shows a schematic of a representative sampled structure in that stretched configuration.

To go further in the structural features of the peptide, we looked at the edge-to-edge distances. We need to check whether the stretched conformation found in the case in which the peptide is part of the whole interface is also maintained when it is extracted from the interface. Indeed, in the ideal case of being introduced as a potential therapeutic, the isolated peptide needs to be in a stretched conformation close enough to that of the dimeric interface to be efficient in properly binding the other chain and neutralising most of the original protein-protein contacts. Here, the edge-to-edge distance is used as a measure of the elongation of the conformation of the peptide [25] and is computed as the distance between the two terminal *Cα*s. The related distributions are illustrated in the right panels of Fig. 2. We first evaluated the extension that the peptide explores during the simulation, where it belongs to the original interface. We therefore chose to use, as a reference, the range of distances over which the peptide spends the vast majority of the simulation time. In particular, we selected the interval as the two standard deviation region centered around the average, accounting for 92% of simulating time (shaded area in the right panel of Fig. 2a). We then referred to this extension when analyzing the edge-to-edge distance in the other two dynamics. As one can see from the distributions on the right of Fig. 2b and Fig. 2c, which show the edge-to-edge distribution for the case in which the peptide is isolated and when it is bound to the other chain, in both cases the peptide spends a significant fraction of the simulating time in the stretched conformation (∼59% and ∼42%, respectively). Hence, despite the expected conformational heterogeneity of the isolated peptide in solution and the occurrence of intra-chain contacts, it still assumes a linear configuration for a considerable amount of time. The absence of durable odd interactions within the peptide structure may also be attributed to the chosen length, as a 15-residue-long peptide is rarely involved in secondary structure organization [36].

Along with these findings, we moved to optimize the identified peptide to enhance its antagonist activity, as discussed in the next section.

### B. Peptide sequence optimisation via computational mutagenesis

Subsequent to the determination of a potentially suitable peptide, we performed the optimisation of the selected sequence in order to make it increasingly competitive for chain binding and, thus, for dimerisation impairment.

The approach used for optimising the peptide is an expansion of a mutagenesis Monte Carlo protocol previously worked out by this team and which has already proven its efficacy in several molecular systems [29] [25] [28] [23].

The preliminary step of this protocol is the calculation of descriptors used to evaluate the complementarity between two interfaces. As mentioned in the introduction, in this work we employed descriptors of shape complementarity, electrostatic complementarity, and hydropathy compatibility (See Methods and Materials section for details about the procedure used to measure these quantities). Since, for a 15-residue sequence with a possible alphabet of 20 amino acids, the space of potential mutants is very large, a Monte Carlo algorithm is employed to efficiently explore this space. According to this strategy, in each step, we introduce a single random mutation in the interface while keeping the other residues unchanged. The mutated structure is predicted using the Scwrl4 software [37] and the chemical-physical descriptors were computed. The evaluation of the introduced mutation is then subjected to the selection procedure based on the Metropolis algorithm working on the difference in terms of the chosen descriptors. In addition to the descriptors, we added a term penalizing excessive mutations to preserve the original peptide sequence, which is already engaged with the target protein. A schematic representation of such a protocol can be found in Fig. 3a.

**FIG. 3:**
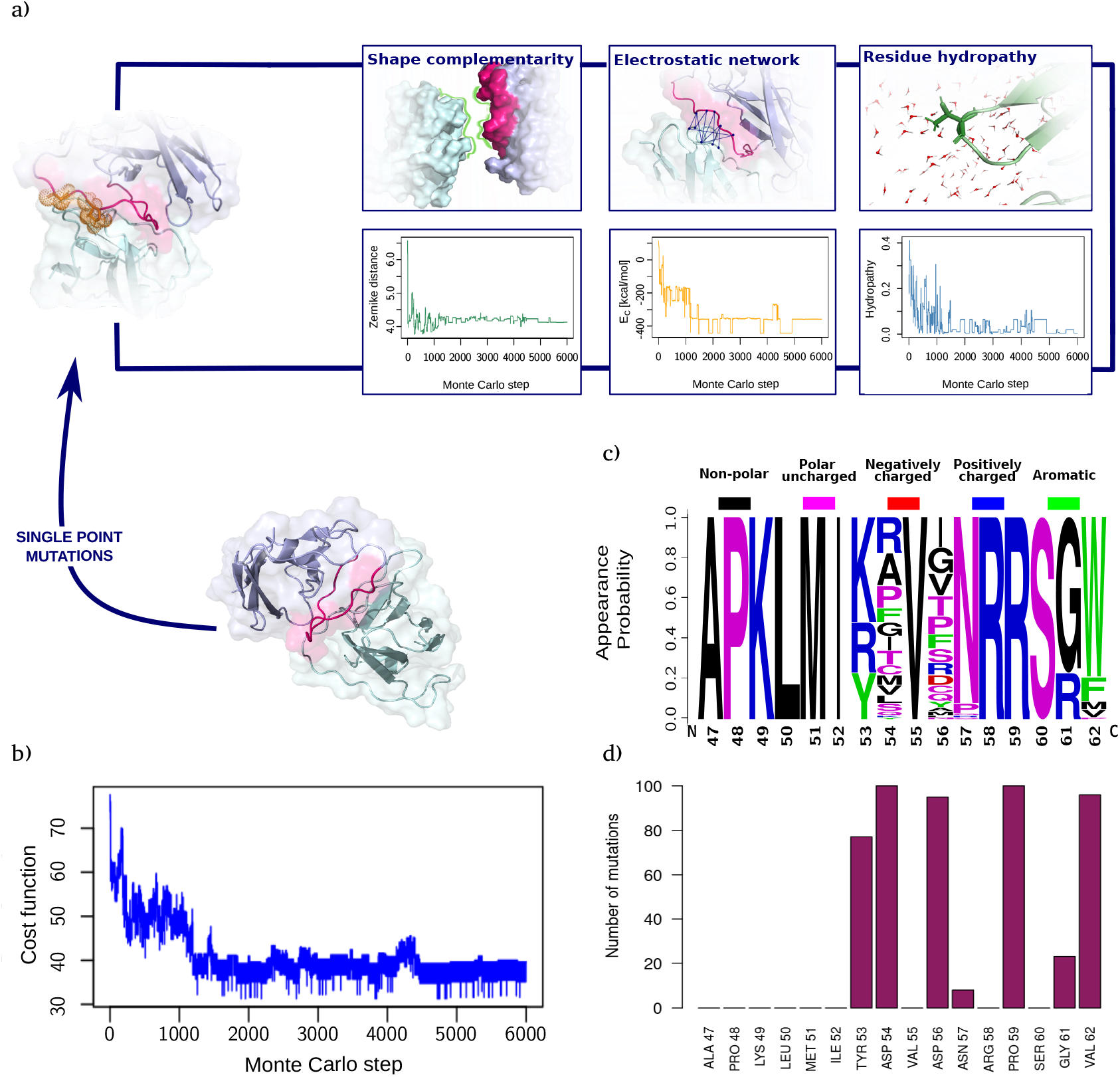
Peptide optimisation through Monte Carlo mutagenesis protocol. **a)** Schematic representation of the main contributions to the Monte Carlo cost function (upper panels) and the respective trends along the 6000 simulation steps of one round over the ten performed (lower panels). **b)** Example of cost function trend for one of the ten MC runs. **c)** Residue appearance probability as provided by WebLogo [38] for the best 100 peptides obtained from the optimisation workflow. The dimension of each letter represents the probability of occurring at a specific position. The physico-chemical properties of the residues (non-polar, polar uncharged, negatively charged, positively charged, aromatic) are highlighted with different colors (respectively: black, magenta, red, blue, green). **d)** Number of mutations involving each residue in the best 100 optimized peptide sequences.

Hence, the cost function (*C*_*f*_) employed in this protocol is:

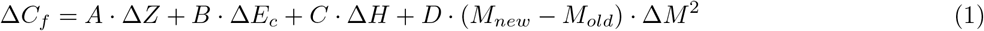

where A, B, C, and D are numerical coefficients chosen to normalize the values of the four terms in this function and to make them equally important. Δ*Z* represents shape complementarity, Δ*E*_*c*_ the Coulomb potential energy, Δ*H* the difference in hydrophobicity between the two interacting regions, and M the number of mutations relative to the original sequence [29].

The acceptance probability is:

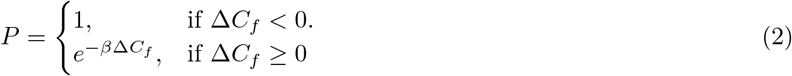

which is a standard for Monte Carlo schemes.

We performed 10 independent Monte Carlo simulations, each consisting of 6,000 steps. Fig. 3b shows the progression of the cost function as a function of simulation steps. Among the 60,000 peptide sequences explored, we filtered out those with more than five mutations relative to the original sequence, selecting the top 100 candidates based on the physicochemical terms of the cost function.

Fig. 3c and d show an analysis of the obtained peptide sequences by depicting the probability of an amino acid to occur at a certain position as provided by the WebLogo webserver [38] (panel c) and the probability of that position to undergo mutations (panel d). According to Fig. 3c, frequently changing residues such as ASP 54 and ASP 56 display very heterogeneous mutability, although they are substituted by amino acids that are not characterized by a net charge. This evidence reflects the necessity of removing a negative charge from those positions, that in both cases originally display an ASP. Oppositely, while PRO 59 is a highly mutating spot as well, it always changes into an arginine thus introducing a positive charge in that position. Moreover, looking at Fig. 3d, it is interesting to note that the mutations are concentrated in very few positions. This ensures that the simulations, although independent, have roughly converged to the same minimum. It is even more remarkable to observe that the most mutated positions are among those that in Fig. 1b are the least involved in the interaction. This result is a first indication of the functionality of the proposed mutagenesis protocol in producing a globally improved structure: indeed, the protocol choose to change the weakest interacting sites while maintaining those that already constitute an optimal combination.

In order to identify a single candidate peptide, among the best 100 *C*_*f*_ s we chose the peptide as the one characterized by the most favorable binding affinity as predicted by the PRODIGY tool [32]. A molecular representation of the newly found peptide compared with its original sequence is presented in Fig. 4a.

**FIG. 4:**
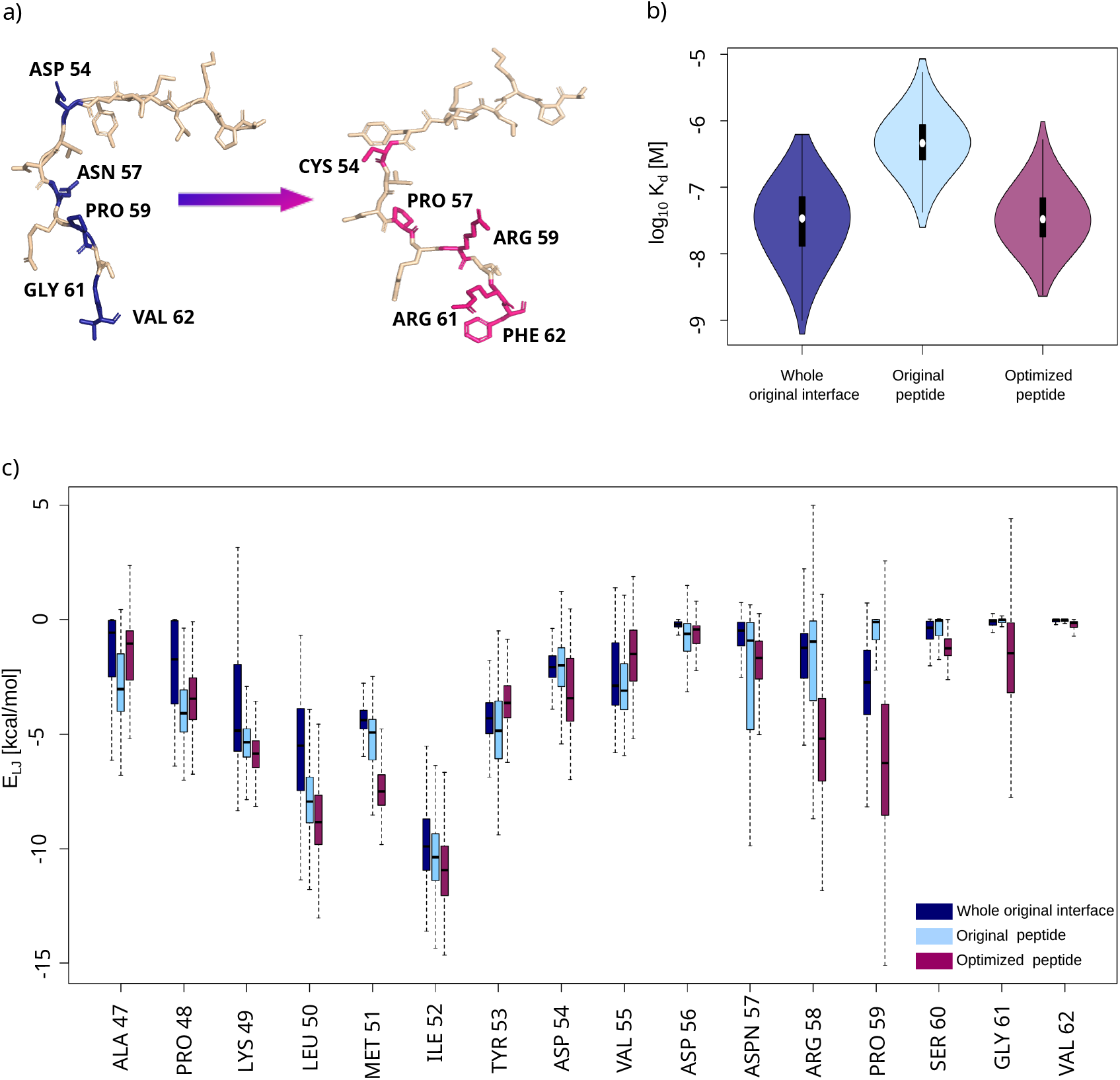
Best optimized peptide validation. **a)** Cartoon of the original peptide sequence and the best optimized sequence obtained. In bold are highlighted the mutated sites and the corresponding substituting residues. **b)** Comparison of the affinities for the pathogenic light chain observed during the dynamics of the whole pathogenic interface (navy), of the original peptide when not surrounded by other interface residues (light blue) and of the optimized peptide (magenta). **c)** Lennard-Jones interaction of each peptide residue in the three cases mentioned in b).

### C. Validation of the optimized sequence

In this section, we assess the ability of the Monte Carlo algorithm to design a peptide capable of inhibiting the interaction between two copies of the misfolded protein, thereby preventing aggregation. In other words, we need to test the competitiveness of the selected peptide with respect to the pathogenic protein-protein binding.

Hence, we performed 1*µ*s-long molecular dynamics simulation of the molecular complex formed by the mutated peptide and its molecular partner. Thus, to obtain an independent validation, we used PRODIGY to predict the binding affinity. In particular, we predicted the binding affinity for each frame extracted from: (i) the dynamics of the whole original pathological dimer, (ii) the dynamics of the complex formed by one monomer and the original peptide, (iii) the dynamics of the complex formed by one monomer and the optimized peptide.

Figure 4b shows a boxplot representation of the affinities obtained in these cases. Inspecting this plot, it is evident that the complex formed by the original peptide and a chain displays binding affinities higher than the range of affinities characterising the entire original interface. This is an expected result since, in the case of the peptide-chain complex, the binding interface size is reduced, thus decreasing the contribution to the complex’s tightness. Besides, the bound optimized peptide spans a way lower range of affinity values than the original peptide-chain complex, thus confirming that our Monte Carlo approach is successful in providing an optimized sequence. Moreover, further important evidence is that the optimized peptide explores conformations whose binding affinity is comparable to that of the dimer, where the whole pathogenic interface is involved.

The improvement introduced by the protocol is also confirmed by the comparison of the vdW interface interaction per residue between the three cases, as depicted in Fig. 4c. Here, we can see that most sites in the optimized peptide share stronger favorable interactions than in the other configurations. This evidence is true not only for those residues that have been mutated (a clear example is PRO 59, where the substitution with a positive charge produces a strong increase in favorable vdW energy), but also for those that have not changed, like ILE 52, which, in the optimized peptide, displays even stronger interactions. Hence, mutations must have repercussions on residues that are not in the closest vicinity. These outcomes confirm that binding enhancement, rather than being ascribed to specific driving spots, should be viewed as a global result of a well-balanced combination of sequence terms and, straightforwardly, structural features [39] that was successfully achieved with the changes introduced by the workflow proposed in this study.

Finally, with all the above considerations, it is possible to conclude that the sampled peptide sequence can be a suitable competitor for binding and could act as an efficient antagonist of pathological dimerisation, thus representing a potential therapeutic.

## III. CONCLUSIONS

The formation of insoluble protein aggregates is a hallmark of many pathological conditions. In this panorama, the atomic details of the earliest steps of aggregation, particularly the formation of pathological dimers that act as seeds for subsequent oligomerization, are often invisible to experimental techniques, making computational approaches essential for their investigation.

In AL amyloidosis, this challenge is further compounded by the patient-specific nature of the aggregating protein, the antibody light chain, further complicating the identification of pharmacological molecules capable of preventing or even reversing the aggregation process. This notwithtanding, the present study aimed at finding a peptide that could serve as a potential drug candidate against pathogenic dimerisation in a specific case of AL amyloidosis.

To this end, we inspected the dynamics of a dimer consisting in two mutant aggregation-prone light chains, whose structure we were able to predict in a previous study. This analysis pinpointed a sequence of 15 residues that proved to be relevant for binding the pathogenic chains both in terms of interface composition and van der Waals interaction involvement. Thus, we refined an already developed Monte Carlo–based protein interface optimisation protocol: we combined shape complementarity, electrostatics, and hydropathy descriptors to optimize competitive peptide binders able to interfere with pathological dimerisation. The optimized peptide revealed an improved binding affinity for the molecular partner compared with the original sequence, achieving values that are comparable with the whole protein-protein complex.

Our results demonstrate the feasibility of designing inhibitory peptides that mimic and block the misfolded interface, preventing the downstream aggregation cascade. While experimental validation remains necessary, this work underscores the potential of computational mutagenesis to identify therapeutic candidates against protein aggregation. Importantly, the strategy is not limited to light-chain amyloidosis but can be generalized to other systems, offering a rational framework for the rapid design of inhibitory peptides and interface-targeting molecules.

## IV. MATERIALS AND METHODS

### A. Inter-Molecular Energy calculations

Inter-molecular non-bonded interaction energies were computed using the parameters obtained from the CHARMM force field [40]. In particular, given two atoms *a*_*l*_ and *a*_*m*_,van der Waals interactions can be calculated as a 12-6 Lennard-Jones potential:

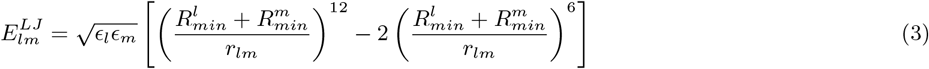

where *r*_*lm*_ is the distance between the two atoms, *ϵ*_*l*_ and *ϵ*_*m*_ are the depths of the potential wells whose minima occur at the distances 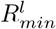 and 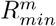, for *a*_*l*_ and *a*_*m*_, respectively.

The total interaction energy between each couple of residues is defined as:

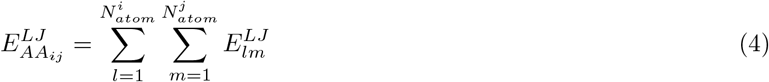

where 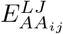 is the energy between two amino acids *i* and *j*, obtained as the sum of the interactions between each atom of the two residues 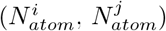.

As for the distance between a pair of residues, this was assessed by selecting the minimum distance between the atoms composing them [41] [42].

### B. Molecular Dynamics parameters

Molecular dynamics simulations were carried out using GROMACS with the CHARMM 27 force field[40]. Energy minimization was performed using the steepest descent algorithm in GROMACS in vacuum, and the procedure was stopped once the maximum force reached 10 kJ/mol [43]. Water was modelled according to the TIP3P three-point model [44]. Before the beginning of the dynamics, the structures were brought to both thermal and pressure equilibrium via Berendsen’s thermostat and barostat and Leap-Frog integrating algorithm.

After the dynamics, a frame at each nanosecond was extracted, so that we ended up with 1000 frames (i.e., structures) for each monomer.

### C. Post-MD analysis

Given a structure, the RMSD measures the average distance between its atoms and those of a reference structure. It can be written as follows:

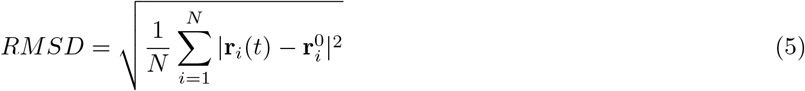

where **r**_*i*_(*t*) is the position vector of the *i*th atom at a given time *t* and 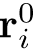 represents the position of the same atom in the reference structure, which in our case corresponds to the configuration prior to the start of the dynamics

### D. Monte Carlo cost function terms

We performed 10 runs of 6000 Monte Carlo steps. The variation of the cost function of the optimisation protocol is given by Eq. 1, where:

– The Δ*Z* term accounts for the variation in shape complementarity between two steps (e.g., *i* + *i* and *i*), assessed as the difference between the Euclidean distance between coefficients of the Zernike polynomial expansion (*Z*_*d*_) of the molecular surfaces:

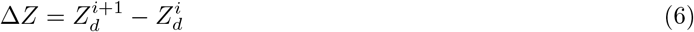
– The term Δ*E*_*c*_ is the variation of the electrostatic potential energy between the two interfaces, computed as a Coulombic potential, in a coarse-grained representation:

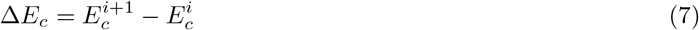 The necessity for leaving an atomistic representation comes from the need to reduce the effects of eventual steric clashes and structural overlaps when mutating a residue.
– The Δ*H* term stands for the change in hydropathy profile that is obtained by employing the scale introduced by Di Rienzo, et al. [24]:

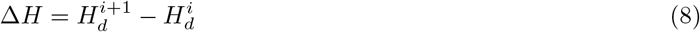

where *H*_*d*_ = *H*^*fixed*^ − *H*^*mut*^ represents the difference between the hydropathic character of the unchanged interface and the mutated one. To assess the hydropathy profile of the two interfaces, the number of occurrences of exposed residues is first counted. The number of such occurrences is multiplied by the hydropathy index associated with those residues. This procedure defines a patch whose mean hydropathy is obtained by dividing the sum of the previously multiplied values by the number of points in the molecular surface.

### E. Monte Carlo cost function coefficients

Since each of the above terms is calculated using descriptors of different nature, and due to the different orders of magnitude they take, it is necessary to set the Monte Carlo coefficients in such a way that the respective terms are comparable to one another within the summation of the cost function. To do so, we set the coefficients *A* = 10, *B* = 0.04, *C* = 32.1, *D* = 0.5.

### F. Computation of molecular surface

The solvent-accessible surface of all protein structures, based on their X-ray coordinates in PDB format [45], was calculated using DMS [46] with a density of 5 points per *Å*^2^ and a water probe radius of 1.4 Å. Unit normal vectors at each surface point were obtained using the -n flag.

The resulting molecular surface consists of a set of points in a three-dimensional Cartesian space (i.e., it is a discretization of the continuous molecular surface). Given a region of interest on this surface, we define a surface patch, Σ, as the points of the surface contained in the region of interest.

### G. 2D Zernike polynomials and invariants

Once the patch is selected, we compute the average of the external normal vectors and reorient Σ in such a way that the average normal vector is aligned with the z-axis. Then, given a point *C* on the z-axis we define the angle *θ* as the largest angle between the perpendicular axis and a secant connecting *C* to any point of the surface Σ. *C* is then set in order that *θ* = 45°. Let us call *r* the distance between *C* and a surface point. We then construct a square grid, and each pixel thus obtained is associated with the mean *r* of the points it contains. In this way, a 2D function representing the patch is retrieved.

Given a function *f* (*r, ϕ*) expressed in polar coordinates and defined within a unitary circle (*r <* 1), it is possible to represent this function in the Zernike basis as

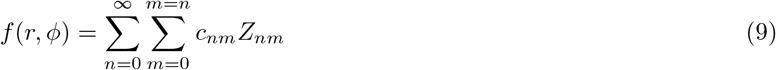

where

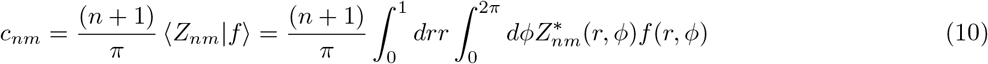

are the expansion coefficients. Zernike polynomials are complex functions, therefore holding a radial and an angular part,

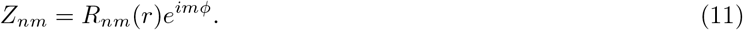

The radial part for a certain couple of indices, *n* and *m*, is given by

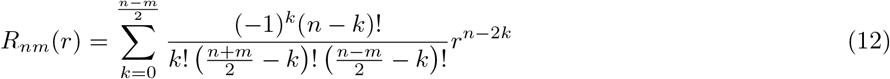

In general, for each couple of polynomials, it can be shown that

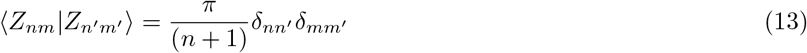

which ensures that the polynomials can form a basis. Knowledge of the set of complex coefficients, {*c*_*nm*_} allows a univocal reconstruction of the original image (with a resolution that depends on the order of expansion, *N* = *max*(*n*)). We found that with *N* = 20, i.e 121 coefficients, a good visual reconstruction of the original image is achieved.

By taking the modulus of each coefficient (*z*_*nm*_ = |*c*_*nm*_|), a set of descriptors can be obtained which have the remarkable feature of being invariant for rotations around the origin of the unitary circle.

The geometric similarity between the two interacting surfaces can then be assessed by comparing the Zernike invariants of their associated 2D projections. In particular, the similarity between patch *i* and *j* is measured as the Euclidean distance between the invariant vectors, i.e.

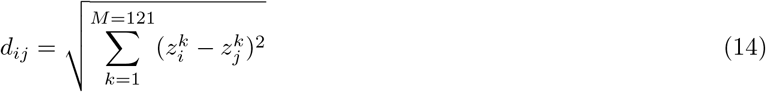

### H. Coarse-graining for electrostatic calculations in the Monte Carlo procedure

Before performing the coarse-graining of the examined proteins, we subject the all-atom structure to the PDB2PQR tool from the APBS suite [47], which provides the partial charges at a desired pH of the composing atoms. Then, a 2-bead coarse-grained representation of the molecule is constructed, where each residue is now modelled as the average position of the backbone atoms and the average position of the side-chain atoms. The global charges of the beads are obtained as the sums of the respective constituting atoms.

## Acknowledgments

This research was partially funded by grants from ERC-2019-Synergy Grant (ASTRA, n. 855923); EIC-2022-PathfinderOpen (ivBM-4PAP, n. 101098989); Project “National Center for Gene Therapy and Drugs based on RNA Technology” (CN00000041) financed by NextGeneration EU PNRR MUR—M4C2—Action 1.4—Call ”Potenziamento strutture di ricerca e creazione di “campioni nazionali di R&S” (CUP J33C22001130001).

The authors would like to thank the Open University Affiliated Research Centre at Istituto Italiano di Tecnologia (ARC@IIT), part of the Open University, Milton Keynes MK76AA, United Kingdom.

